# Procognitive and neurotrophic benefits of α5-GABA-A receptor positive allosteric modulation in a β-amyloid deposition model of Alzheimer’s disease pathology

**DOI:** 10.1101/2022.09.30.510361

**Authors:** Ashley M. Bernardo, Michael Marcotte, Kayla Wong, Dishary Sharmin, Kamal P. Pandey, James M. Cook, Etienne L. Sibille, Thomas D. Prevot

## Abstract

**INTRODUCTION:** Reduced somatostatin (SST) and SST-expressing GABAergic neurons are well-replicated findings in Alzheimer’s disease (AD) and are associated with cognitive deficits. SST cells inhibit pyramidal cell dendrites through α5-GABA-A receptors (α5-GABAA-R). α5-GABAAR positive allosteric modulation (α5-PAM) has procognitive and neurotrophic effects in stress and aging models.

**METHODS:** We tested whether α5-PAM (GL-II-73) could reverse cognitive deficits and neuronal spine loss in early and late stages of β-amyloid deposition in the 5xFAD model (N=48/study; 50% female).

**RESULTS:** Acute or chronic administration of GL-II-73 reversed spatial working memory in 5xFAD mice at 2 and 5 months of age. Chronic GL-II-73 treatment reversed 5xFAD-induced loss of spine density, spine count and dendritic length at both time points, despite β-amyloid accumulation.

**DISCUSSION:** These results demonstrate procognitive and neurotrophic effects of GL-II-73 in early and late stages of Alzheimer-related β-amyloid deposition. This suggests α5-PAM as a novel β-amyloid-independent symptomatic therapeutic approach.

## BACKGROUND

Cognitive deficits (altered executive functions, reduced working memory, impaired spatial memory or daily planning) are reported in patients with dementia and Alzheimer’s disease (AD) [1]. The buildup of β-amyloid plaques leads to neuronal atrophy, neurodegeneration [2, 3] and increased risk for cognitive decline in AD [4]. Monoclonal antibody-based interventions augmenting clearance of β-amyloid plaques were recently approved, providing some level of cognitive deficit relief and better quality of life for patients [5, 6]. Preclinical and clinical studies confirmed reduced β-amyloid load [7, 8], however, correlating amyloid reductions to clinically relevant cognitive outcomes is debated, and it is uncertain whether they have a beneficial impact on brain cells [9-11]. Cholinesterase inhibitors (Donepezil, Galantamine and Rivastigmine) or NMDA receptor antagonist (Memantine) provide modest and temporary relief and do not reverse neuronal atrophy [12]. Novel therapies that are clinically efficacious for cognitive improvement and that promote neuronal health remain needed.

Designing therapies that target pathologies on a continuum between risk states (age, stress, and depression) [13-15] and AD is a promising approach. A commonality between these conditions is the reduced expression of somatostatin (SST) [16]. Reduced SST expression, SST-expressing neuron numbers and co-localization with β-amyloid plaques are well-documented in postmortem brain of AD patients [17-21] and low SST corticospinal fluid level correlate with cognitive impairment in AD [19]. The SST peptide regulates β-amyloid catabolism by engaging neprilysin-catalyzed proteolytic degradation [22]. However, SST is also a marker of a subtype of GABAergic interneurons [23], which inhibit pyramidal cell dendrites [24]. Recent single cell transcriptomic studies confirmed that SST neurons are a vulnerable inhibitory neuron subtype in AD, and showed that the densities of SST neurons was lowest in AD subject with highest levels of AD pathology, and that preserved SST neuron density correlated with cognitive resilience in AD [25].

What could be a potential mechanism linking SST, AD pathologies and cognitive functions? SST cells signal mainly through α5-GABA-A receptors (α5-GABAA-R) on pyramidal cell dendrites [26, 27]. A 25-30% reduction of α5-GABAA-R binding was reported in the hippocampus [28], as well as transcriptional downregulation in postmortem middle temporal gyrus [29] in AD patients compared to age-matched healthy controls. Together, this suggests that a downregulated dendritic inhibitory pathway, comprised of SST cell, α5-GABAA-R and pyramidal cell spines, may contribute to the pathophysiology of AD.

GABA signals through GABAA-Rs, ligand-gated chloride ion channels commonly assembled from two α, two β, and one γ subunit. GABA binds at the interface between α and β subunits of GABAA-Rs, whereas a separate allosteric site lies at the interface between α and γ subunit [30]. Α5-GABAA-Rs are involved in learning and memory functionally and due to their localization in the hippocampus and prefrontal cortex [31-34]. Loss or reduced function of SST-cells or α5-GABAA-R affects cognitive performance through altered information processing by cortical microcircuits, together contributing to cognitive dysfunction [27, 35]. Hence, augmenting α5-GABAA-R function has therapeutic potential for reversing cognitive deficits in conditions characterized by reduced dendritic inhibition, such as stress-related disorders, depression and AD [26, 27]. In support of this hypothesis, we showed that GL-II-73, an α5-GABAA-R preferential positive allosteric modulators (α5-PAM), reversed cognitive deficits in chronic stress and aging models in mice [36]. Moreover, chronic GL-II-73 treatment displayed an unexpected neurotrophic effect, manifested as increased dendritic length and spine density of pyramidal neurons in the prefrontal cortex and hippocampus in stress and aging mouse models [34, 37].

Here, we tested whether α5-PAM (GL-II-73) could reverse cognitive deficits and neuronal spine loss using the 5xFAD mouse model (https://www.alzforum.org/research-models/5xfad-b6sjl). 5xFAD mice constitutively accumulate β-amyloid plaques, starting at 1.5 month of age, with progressive emergence of cognitive deficit and neuronal damage and loss [38, 39]. We predicted that acute and chronic α5-PAM treatment would prevent or reverse cognitive and neuronal deficits in the early stage of β-amyloid deposition (2-3 months of age) and have reduced or no efficacy in late stage (6 months of age) of β-amyloid deposition.

## METHODS

### Ethical Statement

All animal work was performed in accordance with the Ontario Animals for Research Act (RSO 1990, Chapter A.22), Canadian Council on Animal Care (CCAC), and was approved by the Institutes Animal Care Committees (Protocol #846). For study timelines, please refer to **Figures 1A** and **2A**.

**Figure 1.**
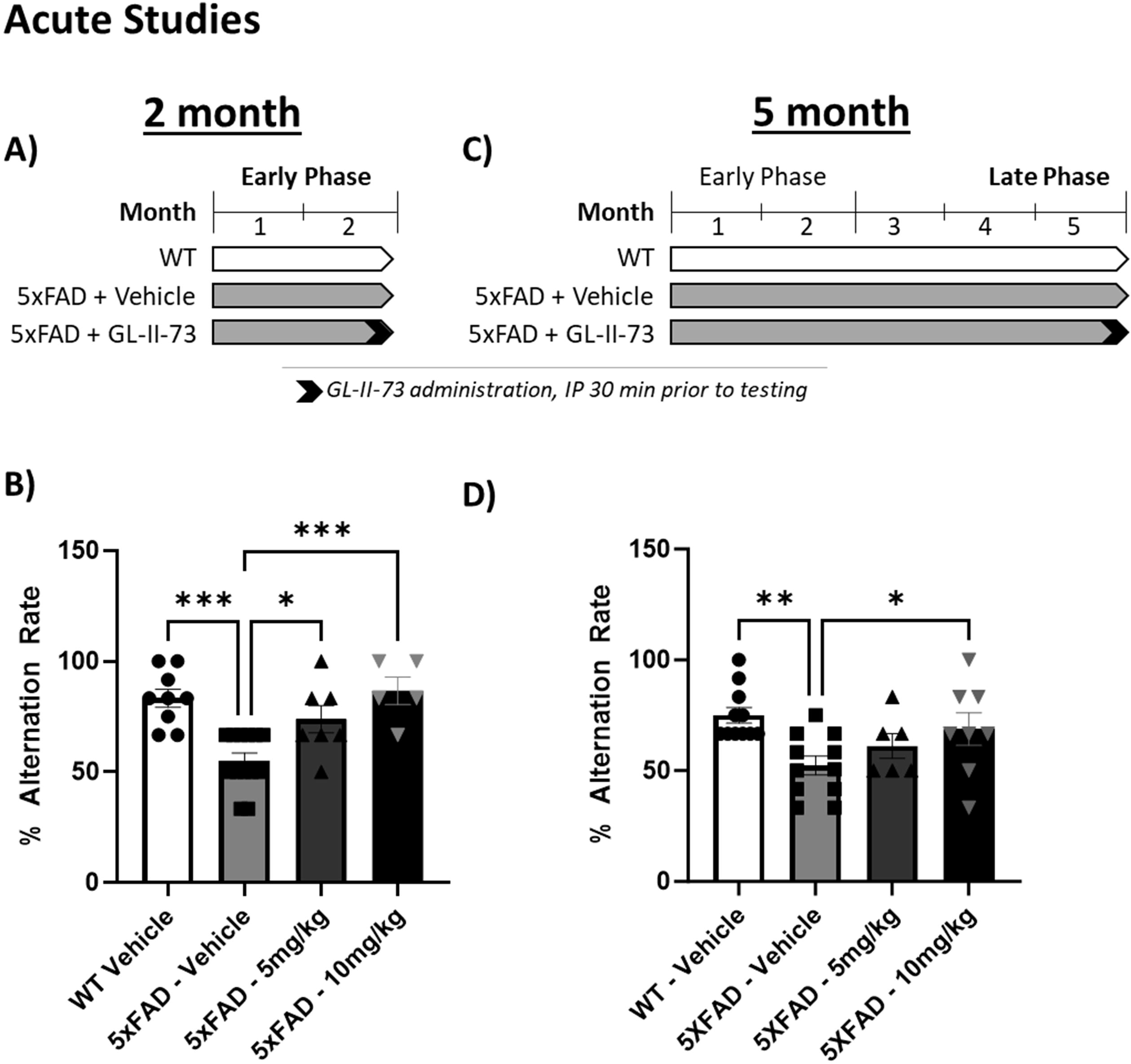
Acute administration of GL-II-73 reverses working memory deficits in 5xFAD mice. 5xFAD and wild type littermate mice were tested in the Y-maze 30 min after an acute administration of GL-II-73 (i.p., 5 or 10 mg/kg). Testing was performed at 2 months (A) or at 5 months (C) of age. 5xFAD mice receiving vehicle displayed significantly reduced alternation rate at 2 (B) and 5 (D) months of age, with 50% alternation rate representing random choice. Administration of GL-II-73 reversed this deficit at 5 and 10 mg/kg in 2 months old 5xFAD mice (B), and at 10 mg/kg in 5 months old 5xFAD mice (D). *p<0.05, **p<0.01, ***p<0.001 compared to 5xFAD-Vehicle.

**Figure 2.**
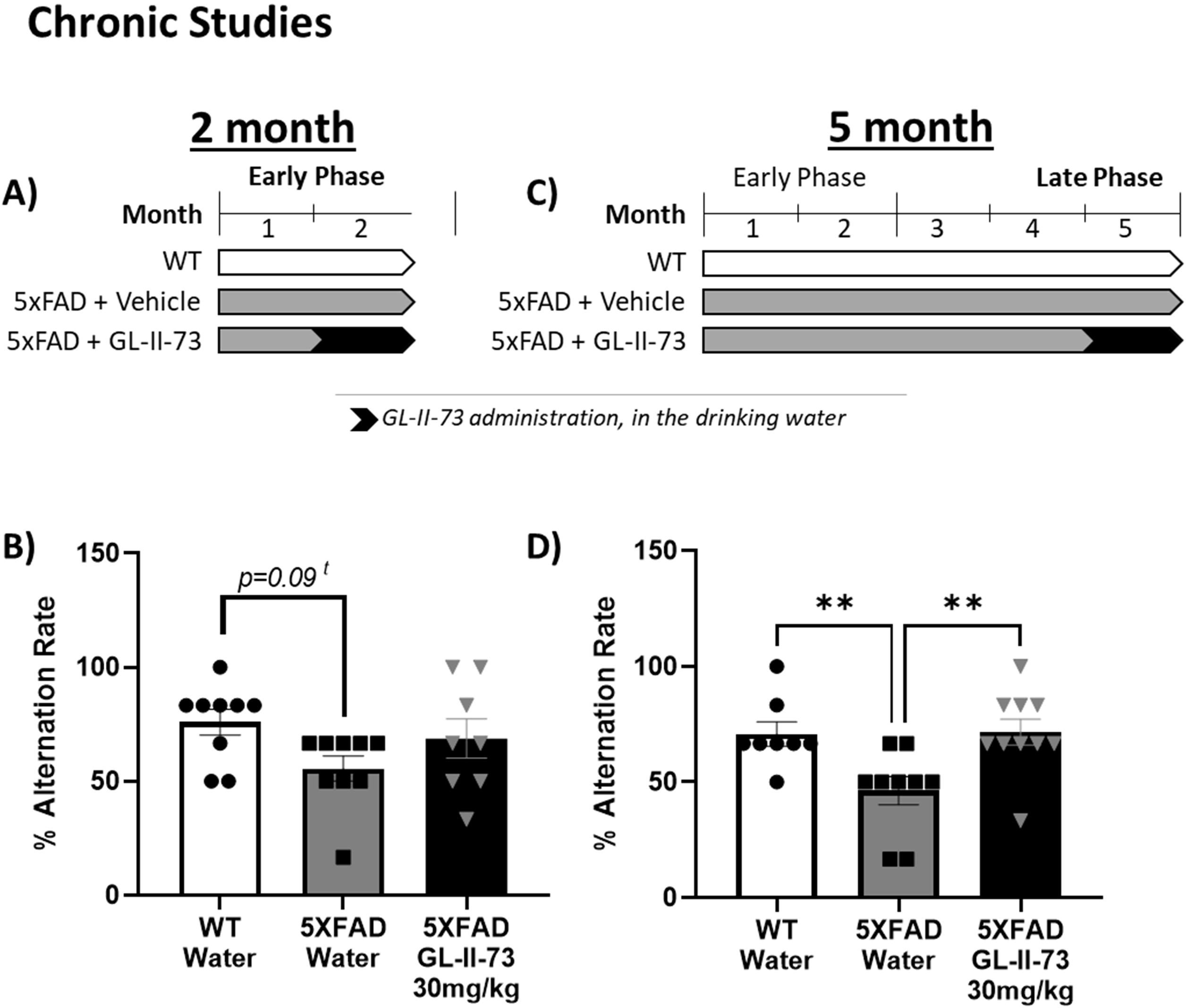
Chronic administration of GL-II-73 reverses working memory deficits in 5xFAD mice. 5xFAD mice received chronic administration of GL-II-73 in their drinking water (30mg/kg) for 4 weeks, prior to being tested in the Y maze at 2 months (A) and 5 months (C) of age. 5xFAD mice receiving water showed a trend in reduced alternation rate at 2 months of age and a significant reduced alternation rate at 5 months of age, which was reversed by chronic GL-II-73 administration. *t=trend p<0.1,* *p<0.05, **p<0.01 compared to 5xFAD-Vehicle.

### Animals

5xFAD breeders were obtained from Jackson Laboratories (Cat #034840-JAX) and paired with WT breeders (Cat #10012) to generate parents of the experimental groups. The following cohorts of 5xFAD heterozygous and WT littermates were generated: Acute studies at 2 month (N=24, 48% female, **Fig. 1A,B**) and 5 month (N=46, 50% female, **Fig. 1C,D**); Chronic studies at 2-3 months (N=48, 50% female, **Fig. 2A,B**) and 5-6 months (N=48, 50% female, **Fig. 2C,D**). Animals were group-housed until the beginning of the study, after which time they were single-housed. Animals had *ad libitum* access to food and water. All animals were maintained on a 12-hour light/dark cycle. Animals were gently handled following the technique described in Marcotte et al [40] to limit their reactivity to the experimenter.

### Drug Preparation and administration

GL-II-73, a compound with preferential activity at the α5-GABAA-R (α5-PAM) [36] was synthesized in Dr. Cook’s group (University of Wisconsin-Milwaukee). For acute administration performed intraperitoneally (IP), GL-II-73 was dissolved in vehicle solution (85% distilled H2O, 14%propylene glycol (Sigma Aldrich), 1% Tween 80 (Sigma Aldrich). The 2 month and 5 month acute studies used GL-II-73 at 5mg/kg and 10mg/kg in the Y maze via IP injection. IP injection with vehicle or GL-II-73 were performed 30 min prior to testing. For chronic studies, GL-II-73 was dissolved in tap water and given to mice in their water for per oral (PO) administration at 30mg/kg, based on calculation of daily water intake.

### Y maze

The Y maze uses spontaneous alternation to investigate spatial working memory. This test was adapted from the T maze paradigm described in Faucher et al [41]. The maze consists of 3 arms (2 goal arms and 1 starting arm) each separated by 120°. The two “goal” arms have a sliding door near the center of the maze and closest to the experimenter is the starting arm with a sliding door at the distal end of the arm that is used as a “start box”. The first phase of the test involves a 2-day habituation phase where mice can freely explore the maze for 10 minutes each day. On the third day, a training phase is conducted. Training consists of placing the animal inside the start box and after a 30 seconds intertrial interval, the sliding door is opened. The mouse can then enter either of the goal arms. Once chosen, the sliding door of the goal arm is closed and the mouse remains in the chosen arm for 30 seconds before being returned to the “start box”. The chosen arm and latency to decide are recorded. This is repeated for 7 trials. On the 4^th^ day a similar procedure is used, however the intertrial interval was increased to 60 seconds and an 8^th^ trial is conducted with a 10 second intertrial interval. This 8^th^ trials evaluates an animal’s motivation to alternate, therefore if no alternation is made, the animal is excluded from the analysis to avoid biases from lack of motivation instead of cognitive performance. Mean % alternation was calculated using Mean alternation %=[(Number of alternations/number of trials)*100]. The maze was wiped with 70% ethanol between trials to remove olfactory cues.

### Golgi Staining and Analysis

At completion of the in-life portion, animals were euthanized via rapid cervical dislocation followed by decapitation. Brains from 6 mice per group (3 males and 3 females), selected pseudo-randomly, based on their behavioral outcomes in the Y maze, to be representative of the full group, were extracted and immediately placed in Golgi Solution for 2 weeks. Brains were then transferred to a shipping solution, coded for blind analysis and sent to NeuroDigitech (San Diego, CA, USA). Brains from the other animals were split into 2 hemispheres (1 for ThioS staining described in the next section, and 1 for molecular analyses not included in this report).

For the brains placed in Golgi staining solution and shipped to NeuroDigiTech, 100µm thick serial, coronal slices were made using a cryostat from anterior to posterior. Slices were mounted on glass slides and the basal and apical dendrites of pyramidal neurons of layers II/III were identified as regions of interest (ROIs) in the PFC and CA1 of the hippocampus. NeuroLucida v10 (Microbrightfield, VT) software was used for quantification on a Dell PC computer and controlled a Zeiss Axioplan 2 image microscope with Optronics MicroFire CCD camera (1600x1200). Motorized X, Y and Z focus enabled high resolution images and later quantification. Low magnification (10x and 20x) was used to identify ROIs with the least truncations of distal dendrites at high magnification (40x-60x). Using immersion oil and Zeiss 100x, 3D images were constructed to allow for spine counting along the complete dendritic tree. Inclusion and exclusion criteria were adapted from Wu et al [42] with inclusion requiring chosen neurons (n=6 per animal) to visually have the dendrites and soma completely filled, no overlapping with other soma and have complete 3D visualization of the dendritic tree. Exclusion criteria was neurons with incomplete staining and/or incomplete visualization. Spine sampling only included those that were well characterized and orthogonal to the dendritic shaft because those above or below are unable to be distinguished well. Raw data was extrapolated and quantified using NeuroExplorer (Microbrightfield, VT, USA).

### Thioflavin S Staining and Analysis

Upon brain extraction, one hemisphere was drop-fixed in 4% PFA for 48 hours then immersed in 30% sucrose for 48 hours. Using a cryostat 30µm serial, coronal slices were taken and stored in cryoprotectant. The free floating Thioflavin S (ThioS) protocol involved 3 washes in PBS for 5 min followed by a 10 min wash in ThioS (1% weight/volume dissolved in ddH2O). 2 washes for were then completed in 75% Ethanol for 5 min each wash followed by rehydration via 3 washes in PBS, each for 5 min. Slices were then mounted on glass slides. Slides were imaged using an Olympus IX73 inverted LED fluorescence microscope and analyzed using Image J software. Image processing involved selecting the ROI (PFC, overall hippocampus), thresholding the image, converting to binary, converting to masks and analyzing particles.

### Statistics

Statistical analyses were performed using GraphPad Prism, from which graphics were made to report mean±SEM. Y maze data was analyzed using One-Way ANOVA followed by Dunnett’s multiple comparisons test. Morphology Golgi stain results were analyzed using a Repeated Measure ANOVA with Tukey’s post hoc testing. Finally, the ThioS amyloid analysis used a Two-Way ANOVA with a post hoc Bonferroni Multiple Comparisons Test to compare the effect of age, and a non-parametric Kruskall-Wallis test followed by Dunn’s tests for between groups comparisons.

## Results

For all statistical values please refer to **Supplementary Tables** referenced.

### Acute GL-II-73 treatment reverses working memory deficits in 5xFAD mice at 2 and 5 months of age

The Y maze was used to assess spatial working memory in 2-month-old 5xFAD mice (corresponding to an early stage of β-amyloid pathology) and of GL-II-73 treatment **(Fig.1 A-B)**. 5xFAD mice receiving vehicle displayed a significant reduced alternation rate compared to WT. The 5 or 10 mg/kg GL-II-73 treatment dose-dependently reversed alternation rate deficits in 5xFAD mice to levels indistinguishable from WT control mice.

At the age of 5 months, similar profiles were observed with a decrease in alternation rate in 5xFAD mice compared to WT, which was reversed by GL-II-73 administered at 10 mg/kg, and intermediate response at 5 mg/kg, i.e. not different from WT controls nor 10mg/kg-treated-5xFAD mice (**Fig. 1 C-D**).

Statistical details are in **Supplementary Table 1**.

### Chronic GL-II-73 treatment reverses working memory deficits in 5xFAD mice at 2 and 5 months of age

We next assessed whether GL-II-73 maintained procognitive efficacy after chronic treatment in the 5xFAD model. Tested at 2 months of age, no significant group differences in alternation rate were found, but a trend was observed for decreased alternation rate in 5xFAD mice receiving water **(Fig.2 A-B)**. GL-II-73 treated 5xFAD mice were not different from the WT control group.

At 5 months of age, analyses of the alternation rate showed a significant difference between groups, marked by significant decrease in 5xFAD receiving water only, compared to WT and a reversal of this deficit in 5xFAD mice receiving 30 mg/kg GL-II-73 in drinking water, which corresponds to the 10mg/kg i.p in the acute study (See methods) (**Fig.2 C-D**).

Statistical details are in **Supplemental Table 1**.

### Chronic GL-II-73 treatment prevents/reverses pyramidal neuron spine loss in 2 and 5-month 5xFAD mice

Morphological changes in PFC and CA1 pyramidal neurons were quantified using Golgi-Cox staining in the chronic studies at 2 and 5 months of age (**Fig. 3**). In the PFC at 2 months of age, spine densities in the apical and basal segments of pyramidal dendrite were significantly reduced in 5xFAD mice receiving water (**Fig. 3G**). Four-week GL-73 treatment resulted in a complete prevention, with spine densities being at similar levels to control mice, and significantly different from 5xFAD mice receiving water only. In the CA1 of the hippocampus, a similar decrease in spine density in the 5xFAD mice receiving water was observed, as well as a reversal/prevention by chronic GL-II-73 administration in 5xFAD mice (**Fig. 3H**). Statistical details are in **Supplementary Table 2**.

**Figure 3.**
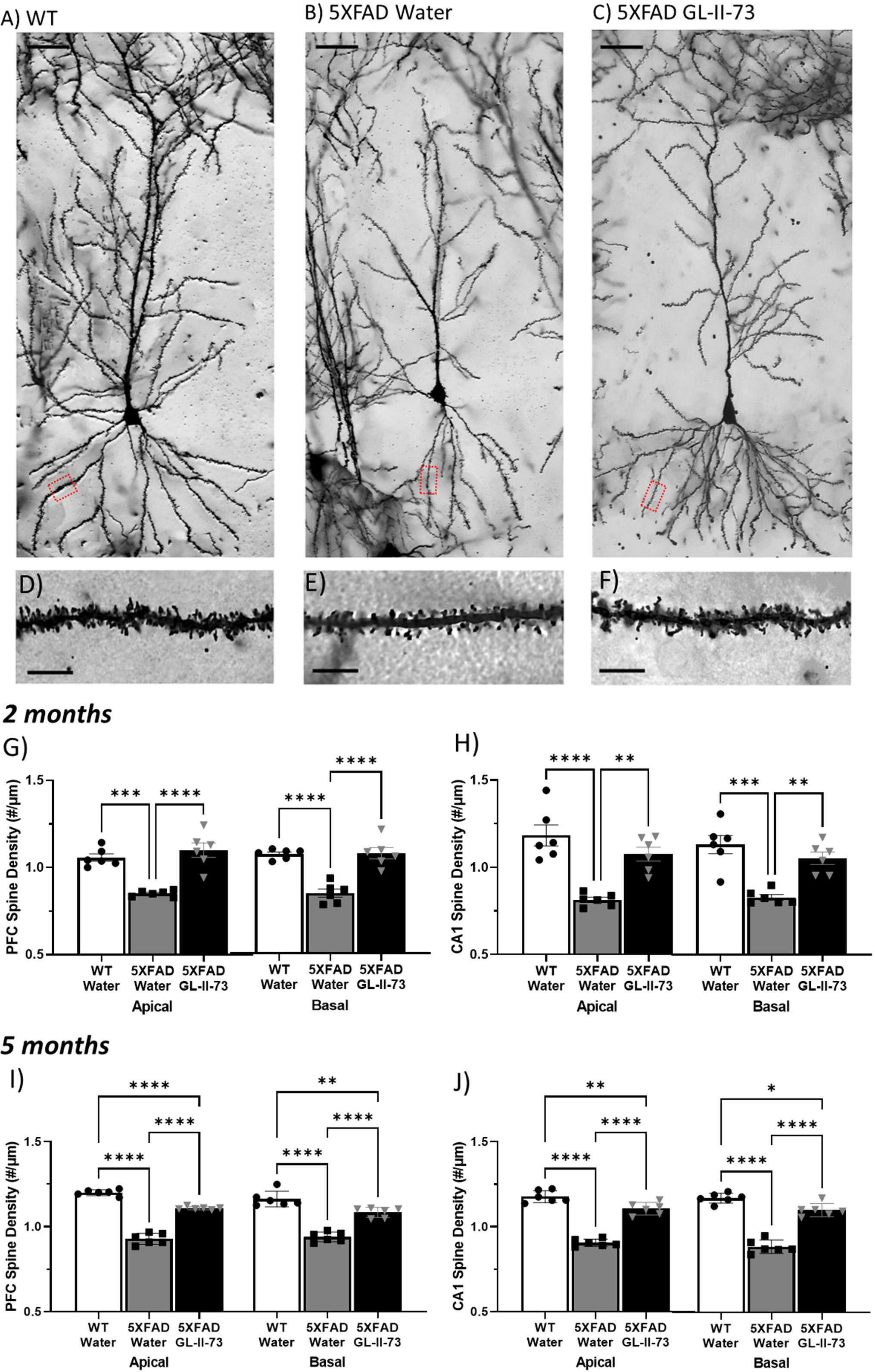
Chronic GL-II-73 reverses 5xFAD-induced morphological changes in pyramidal neurons from the PFC and CA1 of the hippocampus. Using Golgi staining, pyramidal neurons in the PFC and the CA1 were visualized. Representative images of the PFC in 5-month-old chronically treated animals are shown (A-C). Higher magnification representative images of dendrites and spines in the PFC at 5 months are shown (D-F). At 2 months, 5xFAD mice receiving water showed reduced spine density in the apical and basal segments of the pyramidal neurons from both the PFC and the CA1. 4-week GL-II-73 administration reversed such reduction in both segments and in both brain regions (G-H). Spine density levels in the treated group are not different from WT. At 5 months, similar decreases were observed in 5xFAD mice receiving water.4-weeks treatment with GL-II-73 reversed the spine density reduction in the PFC (I) and in the CA1 (J) but levels remained statistically lower compared to WT. *p<0.1,* *p<0.05, **p<0.01, ***p<0.001, ****p<0.0001 compared to 5xFAD-water.

In 5 months old mice, similar profiles were observed with decreases in basal and apical spine densities in the PFC (**Fig. 3I**) and the CA1 of the hippocampus (**Fig. 3J**) of 5xFAD mice. Chronic GL-II-73 resulted in significant recovery of spine loss in both brain regions, although at reduced spine density levels compared to WT **Fig. 3I-J**). Statistical details are in **Supplementary Table 3**.

Analyses of spine counts and dendritic length were also carried out and showed similar results, with spine count reduction and dendritic shrinkage in 5xFAD mice receiving water, and partial or full reversal by chronic GL-II-73 treatment (**Supplementary** Fig. 1-2, and **Supplementary Table 2-3**).

After noting the impact of amyloid load and treatment on spines, we investigated whether such effects were observable on all maturation steps of the spinogenesis (**Fig. 4**). In the PFC, the density of all subtypes of spine was reduced in the 5xFAD mice receiving water in both apical (**Fig.4 A**) and basal dendrites (**Fig.4 B**), with the exception of filopodia in basal dendrites. Chronic GL-II-73 treatment reversed these reductions in almost all spine subtypes in the apical segment (except for stubby spines), and in all subtypes in the basal segment (with a partial recovery in stubby and branched spines) (**Fig.4 A-B**).

**Figure 4.**
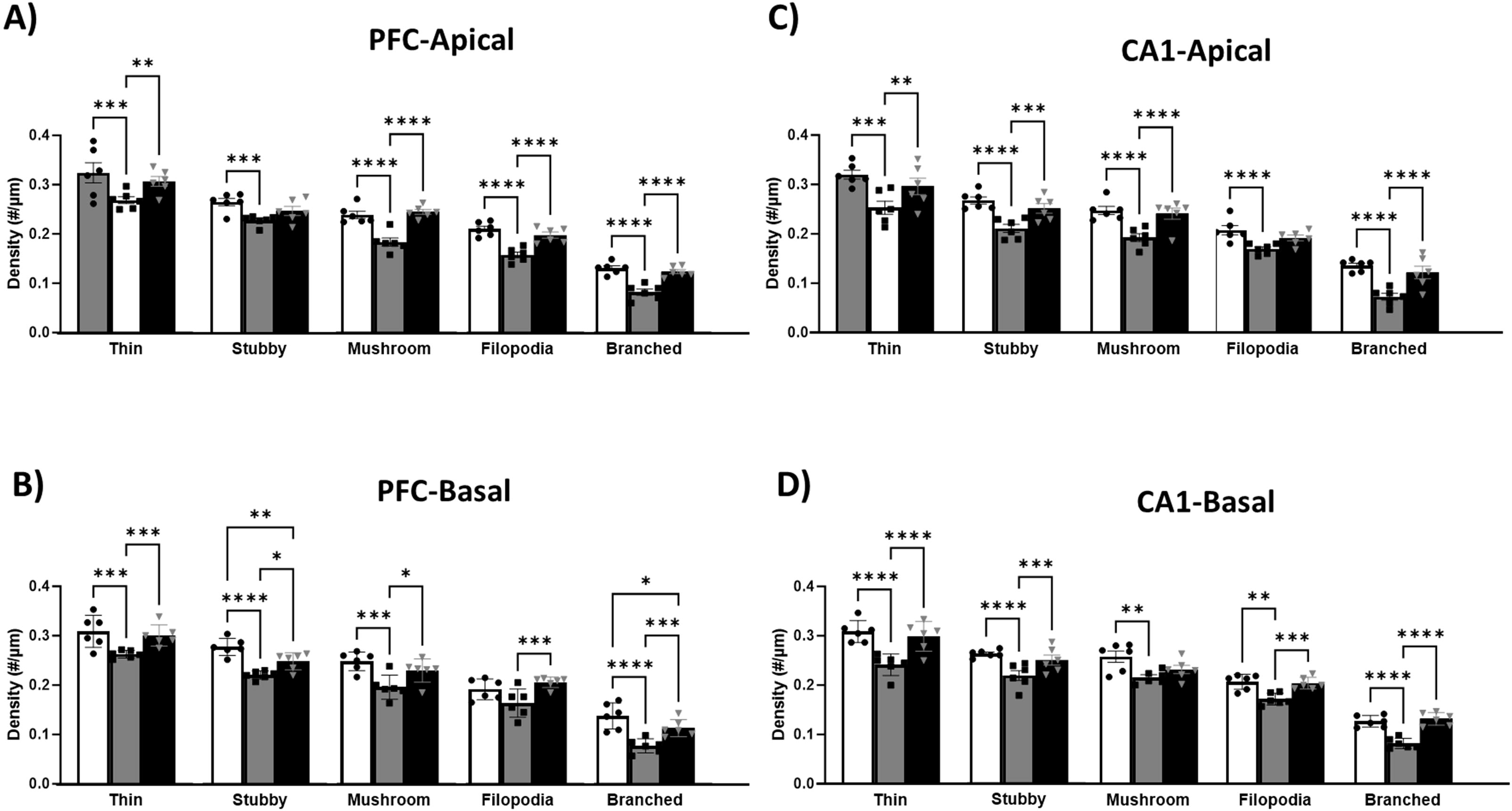
Chronic GL-II-73 administration increases density of all spine maturation steps in pyramidal neurons from the PFC and CA1 of the hippocampus. Presence of all maturation steps of the spines were counted and analyzed in the PFC (A-B) and the CA1 (C-D). 5xFAD mice receiving water exhibited reduced density of all spine subtypes in apical and basal segments of both brain regions (minus the filopodia in the basal segment of dendrites in the PFC that is not significant). 4-week GL-II-73 administration reversed the decrease in most spine subtypes in both brain regions. *P<0.1,* *p<0.05, **p<0.01, ***p<0.001, ****p<0.0001 compared to the 5xFAD-water group.

In the CA1, 5xFAD mice receiving water exhibited reduced density of all spine subtypes in both the apical and basal segments (**Fig.4 C-D**). Chronic GL-II-73 treatment reversed the reduction in density in all spine subtypes, with the exception of filopodia in the apical segment and mushroom in the basal segment (**Fig.4 C-D**). Statistical details are in **Supplementary Table 4**.

### The cognitive and neurotrophic effects of GL-II-73 treatment are independent of β-amyloid load

Β-amyloid load was investigated using ThioS staining. In 5xFAD animals, β-amyloid load increased over time, as expected due to the transgenic construct of the model. 5-month-old 5xFAD mice had higher β-amyloid plaque density in both the PFC and the hippocampus compared to 2-month-old 5xFAD mice, regardless of receiving GL-II-73 or water **(Fig.5 A-N)**. Chronic GL-II-73 had no effect on β-amyloid levels, 2 months 5 months of age in 5xFAD mice. Statistical details are in **Supplementary Table 5 and 6.**

**Figure 5.**
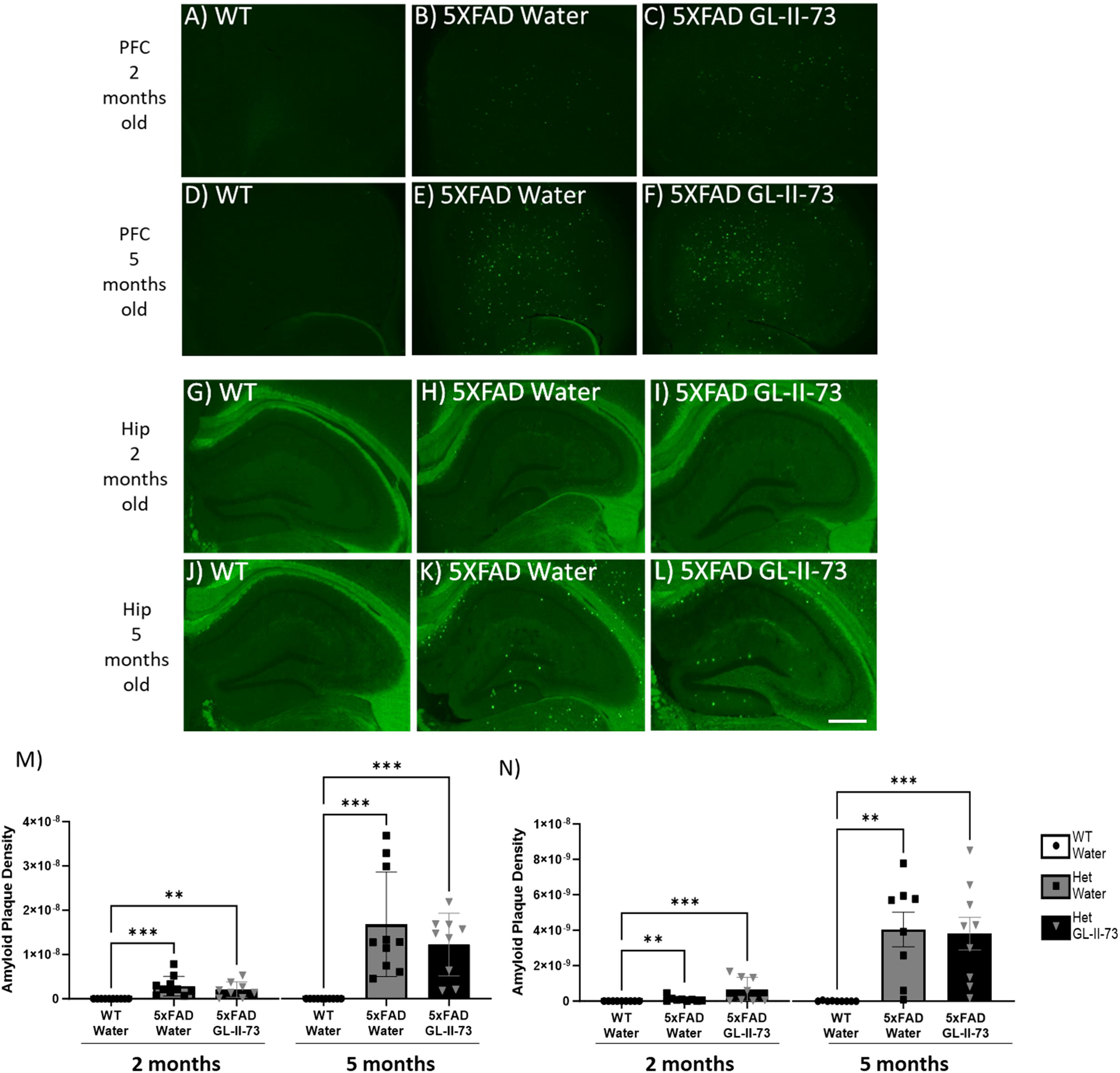
Chronic GL-II-73 administration does not impact β-amyloid load. Representative images of β-amyloid load in the PFC of 2-month (A-C) and 5-month-old animals (D-F) are shown. Representative images of the hippocampus of 2 month old (G-I) and 5month old animals (J-L) are also shown. Age dependent β-amyloid progression was show by a significant increase in total β-amyloid plaque density between 2-month-old 5xFAD water and 5 month old 5xFAD water. A similar finding was found between 2 month old 5xFAD GL-II-73-treated and 5 month old 5xFAD GL-II-73-treated animals. At 5 months of age in both the PFC and the hippocampus, 5xFAD groups showed significantly more β-amyloid. No difference between 5xFAD Water and 5xFAD GL-II-73-treated groups was found. *p<0.05, **p<0.01, ***p<0.001, ****p<0.0001 compared to 5xFAD-water.Scale bar = 400µm

## DISCUSSION

This study investigated the impact of augmenting α5-GABAA-R function, using GL-II-73, an α5-PAM, at early and late stages of β-amyloid deposition on functional, morphological and histological outcomes related to cognition and cellular pathology. The rationale was that the pathway formed by SST-GABAergic interneurons, α5-GABAA-R and pyramidal neuron dendrites is downregulated in AD and risk thereof [43, 44], and that positive α5-GABAA-R modulation has procognitive effects in models of stress [37, 45] and aging [46], known risk factors for AD [34, 37]. We first confirmed that β-amyloid load induced deficits in spatial working memory in 5xFAD mice at 2 and 5 months of ages, corresponding to early and late stages of β-amyloid deposition. We next showed that both acute and chronic GL-II-73 administration reversed these 5xFAD-induced cognitive declines in a dose-dependent manner, including at 5 months of age. We also demonstrated that chronic GL-II-73 treatment reversed dendritic shrinkage and spine loss in 5xFAD mice, affecting all spine maturation steps, at both ages and in both regions investigated, i.e., the PFC and CA1 of the hippocampus, where α5-GABAARs are expressed [47]. As expected due to the transgenic nature of the model, chronic GL-II-73 treatment did not affect β-amyloid accumulation, demonstrating that the procognitive and neurotrophic effects of augmenting α5-GABAA-R function are independent of β-amyloid load.

### Augmenting α5-GABAA-R function reverses spatial working memory deficits in early and late stages of β-amyloid accumulation in 5xFAD mice

We first confirmed the presence of working memory deficits in 5xFAD mice compared to WT at 2 and 5 months of age [39, 48, 49]. This was assessed in the Y-maze test with alternation decreasing from 75-85% to ∼55%. Such reductions are interpreted as working memory deficits since mice do not remember arms they previously visited and instead perform at chance level (∼50%). Interestingly, we report that 5xFAD mice exhibit impairment at earlier times (2-3 month) than the later times (5 months) more frequently investigated [38, 39]. This was fortuitous as we aimed to target an early time of disease progression, for a combined treatment/prevention intervention. As shown by histochemical studies, 2-months of age corresponded to a low but significant β-amyloid load that is sufficient to induce significant spine loss and cognitive deficits. GL-II-73 exhibited a dose-dependent effect in reversing the memory deficit at both ages tested, consistent with prior studies showing dose-dependent effects in chronic model and aging models [34, 37]. Interestingly, the lower dose of 5mg/kg only reversed memory deficits when administered acutely at 2 months of age, not at 5 months, potentially related to lower β-amyloid load at 2 month of age. The effects on working memory were confirmed in the chronic treatment studies, in particular at 5 months of age. In this study, the deficits at 2 months of age was not as strong as in the acute dosing study, consistent with the more variable results reported in the literature at earlier ages, and more robust deficits emerging after 4 months of age in 5xFAD mice [50]. Importantly, 4-week-long GL-II-73 administration robustly reversed this deficit. This sustained efficacy after chronic treatment is consistent with previous studies of chronic GL-II-73 administration in aging [46] or chronic stress [36, 37] models, and confirms the absence of desensitization, as commonly observed with less selective GABAergic drugs [51].

### GL-II-73 has neurotropic effects on pyramidal neurons in the PFC and CA1

Neuronal degeneration in AD is a direct contributor to loss of function, and reduced cognitive performance [52]. The 5xFAD model induces neuronal degeneration, and we confirmed reduced spine count, lower dendritic length and spine density reduction [39, 48, 50] at all levels of spine maturation. Spine pruning, while essential in development [53, 54], could be a contributor to pathological outcomes in neurodegeneration [54]. Mechanisms involved in pruning could be reactivated in neurodegenerative diseases because of toxicity or because of reduced synaptic function [55]. This aligns with the reduced SST function reported in AD [17-21], leading to reduced GABAergic signaling onto pyramidal neurons in the PFC and CA1, ultimately engaging spine pruning. However, mechanisms contributing to this pruning remain underexplored, in particular in the context of AD, and therefore, there is (to our knowledge) no treatment targeting such mechanisms.

Interestingly, GL-II-73 showed reversal of dendritic shrinkage and spine loss in 5xFAD mice in the PFC and hippocampus at the 2- and 5-month time points. Furthermore, spine density was increased at all maturation steps, suggesting a general neurotrophic effect of chronic treatment on spinogenesis. A similar effect on general spine density was reported in the context of aging [46] and chronic stress [37], and is thought to contribute to the sustained effect on working memory, by restoring activity at the synapse, and therefore preventing pruning. We had shown previously that the neurotrophic observed in aging was sustained even 1 week after stopping treatment [46]. How long does this effect last, and whether it also occurs in the context of β-amyloid load in 5xFAD mice remains to be investigated.

Focusing on age of intervention, there was a slight age-dependent effect on the efficacy of GL-II-73 treatment in 5xFAD mice. Indeed, 5xFAD mice receiving GL-II-73 showed spine density level identical to WT mice when treated at 2 months of age. However, the effect of GL-II-73 was less strong at 5 months of age, with spine density levels in GL-II-73-treated mice being different from both non-treated and WT groups, suggesting full recovery at 2 months, but only partial recovery at 5 month of age. This can be due to the increasing β-amyloid load at 5 months of age that could limit the neurotrophic effect of GL-II-73. These mechanisms are currently unknown and the subject of future studies. Previous studies have proposed that GABAergic disinhibition activates AMPARs stimulates TrkB and BDNF neurotrophic pathways [56, 57], which could contribute to the effects observed here. Another point not directly assessed here is whether the effect of GL-II-73 is a reversal (i.e., actively increasing spinogenesis) or a prevention effect (i.e., slowing down neuronal atrophy) The current data suggests that the observed effect is a reversal (at least at the 5-month time point), but studies testing this hypothesis and mechanisms underlying increased spinogenesis are needed.

### The procognitive and neurotrophic effects of augmenting α5-GABAA-R function are independent of β-amyloid load

We confirmed increased β-amyloid density with time/age in the PFC and hippocampus of 5xFAD mice and showed that it is not affected by chronic GL-II-73 treatment. The 5xFAD model uses the constitutively active Thy1 promoter to induce transcription of the transgenes and increase in β-amyloid, which is not subject to endogenous β-amyloid regulation. Hence, a lack of GL-II-73 treatment on β-amyloid load reduction is not surprising, and further shows a lack of effect of treatment on β-amyloid clearance. The notable finding is that, despite an increasing β-amyloid load, we observed improvement in spatial working memory after acute or chronic treatment, as well as neurotrophic effects at both time points. These results suggest beneficial effects of augmenting α5-GABAA-R function, which are independent of β-amyloid load. Considering the newly approved intervention increasing β-amyloid clearance [5, 58], one could anticipate that a combination therapy of an α5-GABAAR PAM with a monoclonal antibody against β-amyloid proteins may represent a promising advancement in the treatment and prevention of AD.

### Limitations

This study focused on working memory. Exploring the impact of GL-II-73 on other cognitive domains impaired in AD could further strengthen the importance of such intervention. Moreover, the behavioral assay and Y-maze apparatus employed here relied on an alternation task that differs from free exploration-based spontaneous alternation task often used in the literature [59, 60], and that is more akin to a T-maze task. Its advantages are a reduced activity and lateralization bias and a more direct testing of working memory and updating function [41]. Another limitation is that the 5xFAD model cannot determine if GL-II-73 treatment impacted plaque deposition due to its constitutive nature. This could be investigated in a model where β-amyloid release is under endogenous promoter control and activity dependent. This is relevant as GABAergic interneurons contribute up to 30% of hippocampal β-amyloid plaque in early stages in the APP model [61]. Finally, AD is not restricted to β-amyloid load. Showing efficacy in additional pathological models of AD, including complex pathophysiology will be important to increase translation potential of a potential α5-PAM therapy.

### Conclusion

Acute and chronic α5-GABAAR potentiation, using GL-II-73, improved spatial working memory deficits in early and late stages of β-amyloid deposition in the 5xFAD model of AD-related β-amyloid load pathology, and produced positive neurotrophic effects in PFC and CA1 pyramidal neurons. The combined data suggests therapeutic relevance for positive modulation of α5-GABAAR function for diseases such as AD.

## ACKNOWLEDGEMENTS

Authors thank Mehrab Ali for support with administrative tasks throughout the study, the CAMH animal facility staff for the caring for our animals over the study duration, the members of NeuroDigitech for their contribution to data generation, and all the animals that contributed to these studies.

## Consent Statement

There was no human participant in this study. Therefore, consent was not necessary

