## Supplementary Material for "Procognitive and neurotrophic benefits of α5-GABA-A receptor positive allosteric modulation in a β-amyloid deposition model of Alzheimer’s disease pathology"

**Positive allosteric modulation of α5-GABA A Receptor in the 5XFAD mouse model has cognitive and neurotrophic benefits**

Ashley M. Bernardo^1^, Michael Marcotte^1^, Kayla Wong^1^, Dishary Sharmin^2^, Kamal P. Pandey^2^, James M. Cook^2^, Etienne L. Sibille^1, 3, 4*^, Thomas D. Prevot^1, 3*^

^1^Campbell Family Mental Health Research Institute of CAMH, 250 college street, Toronto, ON, M5T 1R8 Canada

^2^Department of Chemistry and Biochemistry, University of Wisconsin–Milwaukee, 3210 N Cramer Street, 53211, WI, USA

^3^Department of Psychiatry, University of Toronto, 250 college street, Toronto, ON, M5T 1R8 Canada

^4^Department of Pharmacology and Toxicology, University of Toronto, Medical Sciences Building, 1 King's College Cir Room 4207, Toronto, ON, M5S 1A8, Canada

*Corresponding Authors:

Thomas D. Prevot, Ph.D., CAMH, 250 College Street, room 131, Toronto, ON M5T 1R8, Canada

Etienne Sibille, Ph.D, CAMH, 250 College Street, room 134, Toronto, ON M5T 1R8, Canada

**Supplemental Material**

### ***Supplementary Tables***

**Table 1.** Y maze Behavioral Statistics

| Study | Assay | Statistical Test | F-Value | P-Value | Sex effect |
| --- | --- | --- | --- | --- | --- |
| ***Acute Dosing*** | | | | | |
| 2 Month Acute | Y maze | One-Way ANOVA | F_(3,30)_= 10.78 | **p<0.0001***** | p>0.05 |
|  |  | *Dunnett’s* | *WT vs 5xFAD-vehicle*  *WT vs 5xFAD-5mg/kg*  *WT vs 5xFAD-10mg/kg*  *5xFAD-vehicle vs 5xFAD+5mg/kg*  *5xFAD-vehicle vs 5xFAD+10mg/kg* | **p=0.0001*****  p>0.1  p>0.1  **p=0.017***  **p=0.0003***** |  |
| 5 Month Acute | Y maze | One-Way ANOVA | F_(3,32)_=4.472 | **p=0.01**** | p>0.05 |
|  |  | *Fisher’s LSD* | *WT vs 5XFAD*  *WT vs 5XFAD+GL-II-73 5mg/kg*  *WT vs 5XFAD+GL-II-73 10mg/kg*  *5XFAD vs 5XFAD+GL-II-73 5mg/kg*  *5XFAD vs 5XFAD+GL-II-73 10mg/kg* | ***p=0.001*****  *p=0.08^t^*  *p=0.38*  *p=0.26*  ***p=0.03**** |  |
| ***Chronic Dosing*** | | | | | |
| 2 Month Chronic | Y maze | One-Way ANOVA | F_(2,23)_= 2.553 | p=0.09^t^ | p>0.05 |
| 5 Month Chronic | Y maze | One-Way ANOVA | F_(2,24)_=6.445 | **p=0.0057**** | p>0.05 |
|  |  | *Fisher’s LSD* | *WT vs 5XFAD*  *WT vs 5XFAD+GL-II-73*  *5XFAD vs 5XFAD+GL-II-73* | ***p=0.0068*****  *p=0.9189^t^*  ***p=0.0035***** |  |

**Table 2.** 2 month Morphology Studies

| Region | Assay | Statistical Test | F-Value | P-Value |
| --- | --- | --- | --- | --- |
| PFC | Golgi (Spine density – basal) | Repeated measure ANOVA | F_(2,30)_= 29.55 | **p<0.0001****** |
|  |  | Tukey’s | WT vs 5XFAD  WT vs 5XFAD + GL-II-73  5XFAD vs 5XFAD + GL-II-73 | **p<0.0001******  p=0.99  **p<0.0001****** |
|  | Golgi (Spine density – apical) | Repeated measure ANOVA | F_(2,15)_= 24.10 | **p<0.0001****** |
|  |  | Tukey’s | WT vs 5XFAD  WT vs 5XFAD + GL-II-73  5XFAD vs 5XFAD + GL-II-73 | **p<0.0002*****  p=0.51  **p<0.0001****** |
|  | Golgi (Spine count – basal) | Repeated measure ANOVA | F_(2,15)_= 21.21 | **p<0.0001****** |
|  |  | Tukey’s | WT vs 5XFAD  WT vs 5XFAD + GL-II-73  5XFAD vs 5XFAD + GL-II-73 | **p=0.0004*****  p=0.63  **p<0.0001****** |
|  | Golgi (Spine count – apical) | Repeated measure ANOVA | F_(2,15)_= 12.23 | **p=0.0007***** |
|  |  | Tukey’s | WT vs 5XFAD  WT vs 5XFAD + GL-II-73  5XFAD vs 5XFAD + GL-II-73 | **p=0.02***  p=0.26  **p=0.0006***** |
|  | Golgi (Dendritic Length – basal) | Repeated measure ANOVA | F_(2,15)_= 7.462 | **p=0.006**** |
|  |  | Tukey’s | WT vs 5XFAD  WT vs 5XFAD + GL-II-73  5XFAD vs 5XFAD + GL-II-73 | **p=0.05***  p=0.52  **p=0.005**** |
|  | Golgi (Dendritic Length – apical) | Repeated measure ANOVA | F_(2,15)_= 6.28 | **p=0.01**** |
|  |  | Tukey’s | WT vs 5XFAD  WT vs 5XFAD + GL-II-73  5XFAD vs 5XFAD + GL-II-73 | p=0.17  p=0.27  **p=0.0008**** |
| CA1 | Golgi (Spine density – basal) | Repeated measure ANOVA | F_(2,15)_= 17.47 | **p=0.0001***** |
|  |  | Tukey’s | WT vs 5XFAD  WT vs 5XFAD + GL-II-73  5XFAD vs 5XFAD + GL-II-73 | **p=0.0001*****  p=0.33  **p=0.002**** |
|  | Golgi (Spine density – apical) | Repeated measure ANOVA | F_(2,15)_= 19.6 | **p<0.0001****** |
|  |  | Tukey’s | WT vs 5XFAD  WT vs 5XFAD + GL-II-73  5XFAD vs 5XFAD + GL-II-73 | **p<0.0001******  p=0.21  **p=0.0017**** |
|  | Golgi (Spine count – basal) | Repeated measure ANOVA | F_(2,15)_= 11.38 | **p=0.001**** |
|  |  | Tukey’s | WT vs 5XFAD  WT vs 5XFAD + GL-II-73  5XFAD vs 5XFAD + GL-II-73 | **p=0.0009*****  p=0.42  **p=0.012*** |
|  | Golgi (Spine count – apical) | Repeated measure ANOVA | F_(2,15)_= 15.71 | **p=0.0002***** |
|  |  | Tukey’s | WT vs 5XFAD  WT vs 5XFAD + GL-II-73  5XFAD vs 5XFAD + GL-II-73 | **p=0.0002*****  p=0.19  **p=0.006**** |
|  | Golgi (Dendritic Length – basal) | Repeated measure ANOVA | F_(2,15)_= 4.64 | **p=0.03*** |
|  |  | Tukey’s | WT vs 5XFAD  WT vs 5XFAD + GL-II-73  5XFAD vs 5XFAD + GL-II-73 | **p=0.03***  p=0.72  p=0.11 |
|  | Golgi (Dendritic Length – apical) | Repeated measure ANOVA | F_(2,15)_= 6.715 | **p=0.008**** |
|  |  | Tukey’s | WT vs 5XFAD  WT vs 5XFAD + GL-II-73  5XFAD vs 5XFAD + GL-II-73 | **p=0.007****  p=0.43  p=0.08^t^ |

**Table 3.** 5 month Morphology Studies

| Region | Assay | Statistical Test | F-Value | P-Value |
| --- | --- | --- | --- | --- |
| PFC | Golgi (Spine density – basal) | Repeated measure ANOVA | F_(2,15)_= 61.54 | **p<0.0001****** |
|  |  | Tukey’s | WT vs 5XFAD  WT vs 5XFAD + GL-II-73  5XFAD vs 5XFAD + GL-II-73 | **p<0.0001******  **p=0.004****  **p<0.0001****** |
|  | Golgi (Spine density – apical) | Repeated measure ANOVA | F_(2,15)_= 240.4 | **p<0.0001****** |
|  |  | Tukey’s | WT vs 5XFAD  WT vs 5XFAD + GL-II-73  5XFAD vs 5XFAD + GL-II-73 | **p<0.0002******  **p<0.0001****p<0.0001****** |
|  | Golgi (Spine count – basal) | Repeated measure ANOVA | F_(2,15)_= 42.58 | **p<0.0001****** |
|  |  | Tukey’s | WT vs 5XFAD  WT vs 5XFAD + GL-II-73  5XFAD vs 5XFAD + GL-II-73 | **p<0.0001******p=0.07^t^  **p<0.0001****** |
|  | Golgi (Spine count – apical) | Repeated measure ANOVA | F_(2,15)_= 48.76 | **p<0.0001****** |
|  |  | Tukey’s | WT vs 5XFAD  WT vs 5XFAD + GL-II-73  5XFAD vs 5XFAD + GL-II-73 | **p<0.0001******  **p=0.05***  **p<0.0001****** |
|  | Golgi (Dendritic Length – basal) | Repeated measure ANOVA | F_(2,15)_= 18.55 | **p<0.0001****** |
|  |  | Tukey’s | WT vs 5XFAD  WT vs 5XFAD + GL-II-73  5XFAD vs 5XFAD + GL-II-73 | **p=0.0001*****  p=0.56  **p=0.0008***** |
|  | Golgi (Dendritic Length – apical) | Repeated measure ANOVA | F_(2,15)_= 32.34 | **p<0.0001****** |
|  |  | Tukey’s | WT vs 5XFAD  WT vs 5XFAD + GL-II-73  5XFAD vs 5XFAD + GL-II-73 | **p<0.0001**** p=0.02***  **p=0.0006***** |
| CA1 | Golgi (Spine density – basal) | Repeated measure ANOVA | F_(2,15)_= 104.0 | **p=0.0001***** |
|  |  | Tukey’s | WT vs 5XFAD  WT vs 5XFAD + GL-II-73  5XFAD vs 5XFAD + GL-II-73 | **p<0.0001******  **p=0.0102***  **p<0.0001****** |
|  | Golgi (Spine density – apical) | Repeated measure ANOVA | F_(2,15)_= 118.1 | **p<0.0001****** |
|  |  | Tukey’s | WT vs 5XFAD  WT vs 5XFAD + GL-II-73  5XFAD vs 5XFAD + GL-II-73 | **p<0.0001******  **p=0.004****  **p<0.0001****** |
|  | Golgi (Spine count – basal) | Repeated measure ANOVA | F_(2,15)_= 32.34 | **p<0.0001****** |
|  |  | Tukey’s | WT vs 5XFAD  WT vs 5XFAD + GL-II-73  5XFAD vs 5XFAD + GL-II-73 | **p<0.0001******  **p=0.02***  **p=0.0006***** |
|  | Golgi (Spine count – apical) | Repeated measure ANOVA | F_(2,15)_= 35.82 | **p<0.0001****** |
|  |  | Tukey’s | WT vs 5XFAD  WT vs 5XFAD + GL-II-73  5XFAD vs 5XFAD + GL-II-73 | **p<0.0001******p=0.39  **p<0.0001****** |
|  | Golgi (Dendritic Length – basal) | Repeated measure ANOVA | F_(2,15)_= 30.5 | **p<0.0001****** |
|  |  | Tukey’s | WT vs 5XFAD  WT vs 5XFAD + GL-II-73  5XFAD vs 5XFAD + GL-II-73 | **p<0.0001******p=0.60  **p<0.0001****** |
|  | Golgi (Dendritic Length – apical) | Repeated measure ANOVA | F_(2,15)_= 17.67 | **p<0.0001****** |
|  |  | Tukey’s | WT vs 5XFAD  WT vs 5XFAD + GL-II-73  5XFAD vs 5XFAD + GL-II-73 | **p=0.0002*****  p=0.92  **p=0.0005***** |

**Table 4**. 5 month Morphology Studies - Spine Subtypes

| Region | Subtype | Statistical Test | F-Value | P-Value |
| --- | --- | --- | --- | --- |
| **PFC** | Thin (Apical) | Repeated measure ANOVA | F _(2, 30)_ = 12.08 | **p<0.0001****** |
|  |  | Tukey’s | WT vs 5XFAD  WT vs 5XFAD + GL-II-73  5XFAD vs 5XFAD + GL-II-73 | **P=0.0007*****  p=0.43  **p=0.0024**** |
|  | Stubby (Apical) | Repeated measure ANOVA | F _(2, 30)_ = 6.92 | **P=0.0022**** |
|  |  | Tukey’s | WT vs 5XFAD  WT vs 5XFAD + GL-II-73  5XFAD vs 5XFAD + GL-II-73 | **P=0.001*****  P=0.22  P=0.10 |
|  | Mushroom (Apical) | Repeated measure ANOVA | F _(2, 30)_ = 21.30 | **p<0.0001****** |
|  |  | Tukey’s | WT vs 5XFAD  WT vs 5XFAD + GL-II-73  5XFAD vs 5XFAD + GL-II-73 | **p<0.0001******p=0.82  **p<0.0001****** |
|  | Filopodia (Apical) | Repeated measure ANOVA | F _(2, 30)_ = 27.06 | **p<0.0001****** |
|  |  | Tukey’s | WT vs 5XFAD  WT vs 5XFAD + GL-II-73  5XFAD vs 5XFAD + GL-II-73 | **p<0.0001******  p=0.21  **p<0.0001****** |
|  | Branched (Apical) | Repeated measure ANOVA | F _(2, 30)_ = 22.38 | **p<0.0001****** |
|  |  | Tukey’s | WT vs 5XFAD  WT vs 5XFAD + GL-II-73  5XFAD vs 5XFAD + GL-II-73 | **p=0.0001*****  p=0.61  **p=0.0008***** |
|  | Thin (Basal) | Repeated measure ANOVA | F _(2, 30)_ = 14.88 | **p<0.0001****** |
|  |  | Tukey’s | WT vs 5XFAD  WT vs 5XFAD + GL-II-73  5XFAD vs 5XFAD + GL-II-73 | **P=0.0002***** p=0.67  **p=0.0002***** |
|  | Stubby (Apical) | Repeated measure ANOVA | F _(2, 30)_ = 20.04 | **P<0.0001****** |
|  |  | Tukey’s | WT vs 5XFAD  WT vs 5XFAD + GL-II-73  5XFAD vs 5XFAD + GL-II-73 | **P<0.0001******  **P=0.0048****  **P=0.018*** |
|  | Mushroon (Apical) | Repeated measure ANOVA | F _(2, 30)_ = 15.05 | **P<0.0001****** |
|  |  | Tukey’s | WT vs 5XFAD  WT vs 5XFAD + GL-II-73  5XFAD vs 5XFAD + GL-II-73 | **P=0.0002*****  P=0.12  **P=0.01*** |
|  | Filopodia (Apical) | Repeated measure ANOVA | F _(2, 30)_ = 8.394 | **P=0.0019**** |
|  |  | Tukey’s | WT vs 5XFAD  WT vs 5XFAD + GL-II-73  5XFAD vs 5XFAD + GL-II-73 | P=0.09  P=0.248  **P=0.0005***** |
|  | Branched (Apical) | Repeated measure ANOVA | F _(2, 30)_ = 35.22 | **P<0.0001****** |
|  |  | Tukey’s | WT vs 5XFAD  WT vs 5XFAD + GL-II-73  5XFAD vs 5XFAD + GL-II-73 | **P<0.0001******  **P=0.013***  **P=0.0001***** |
| **CA1** | Thin (Apical) | Repeated measure ANOVA | F _(2, 30)_ = 11.32 | **P=0.0001***** |
|  |  | Tukey’s | WT vs 5XFAD  WT vs 5XFAD + GL-II-73  5XFAD vs 5XFAD + GL-II-73 | **P=0.0008*****  P=0.24  **P=0.007**** |
|  | Stubby (Apical) | Repeated measure ANOVA | F _(2, 30)_ = 18.07 | **P<0.0001****** |
|  |  | Tukey’s | WT vs 5XFAD  WT vs 5XFAD + GL-II-73  5XFAD vs 5XFAD + GL-II-73 | **P<0.0001******  P=0.25  **P=0.0008***** |
|  | Mushroom (Apical) | Repeated measure ANOVA | F _(2, 30)_ =18.69 | **P<0.0001****** |
|  |  | Tukey’s | WT vs 5XFAD  WT vs 5XFAD + GL-II-73  5XFAD vs 5XFAD + GL-II-73 | **P<0.0001******  P=0.81  **P<0.0001****** |
|  | Filopodia (Apical) | Repeated measure ANOVA | F _(2, 30)_ = 10.27 | **P=0.0003***** |
|  |  | Tukey’s | WT vs 5XFAD  WT vs 5XFAD + GL-II-73  5XFAD vs 5XFAD + GL-II-73 | **P<0.0001******  P=0.22  P=0.052 |
|  | Branched (Apical) | Repeated measure ANOVA | F _(2, 30)_ = 38.14 | **P<0.0001****** |
|  |  | Tukey’s | WT vs 5XFAD  WT vs 5XFAD + GL-II-73  5XFAD vs 5XFAD + GL-II-73 | **P<0.0001******  P=0.24  **P<0.0001****** |
|  | Thin (Basal) | Repeated measure ANOVA | F _(2, 30)_ = 20.06 | **P<0.0001****** |
|  |  | Tukey’s | WT vs 5XFAD  WT vs 5XFAD + GL-II-73  5XFAD vs 5XFAD + GL-II-73 | **P<0.0001******  P=0.69  **P<0.0001****** |
|  | Stubby (Apical) | Repeated measure ANOVA | F _(2, 30)_ = 17.41 | **P<0.0001****** |
|  |  | Tukey’s | WT vs 5XFAD  WT vs 5XFAD + GL-II-73  5XFAD vs 5XFAD + GL-II-73 | **P<0.0001******  P=0.26  **P=0.0002***** |
|  | Mushroon (Apical) | Repeated measure ANOVA | F _(2, 30)_ = 7.167 | **P=0.0017**** |
|  |  | Tukey’s | WT vs 5XFAD  WT vs 5XFAD + GL-II-73  5XFAD vs 5XFAD + GL-II-73 | **P=0.002****  P=0.06  P=0.30 |
|  | Filopodia (Apical) | Repeated measure ANOVA | F _(2, 30)_ = 10.11 | **P=0.0003***** |
|  |  | Tukey’s | WT vs 5XFAD  WT vs 5XFAD + GL-II-73  5XFAD vs 5XFAD + GL-II-73 | **P=0.002****  P=0.94  **P=0.0003***** |
|  | Branched (Apical) | Repeated measure ANOVA | F _(2, 30)_ = 20.54 | **P<0.0001****** |
|  |  | Tukey’s | WT vs 5XFAD  WT vs 5XFAD + GL-II-73  5XFAD vs 5XFAD + GL-II-73 | **P<0.0001******  P=0.81  **P<0.0001****** |

**Table 5.** Thioflavin S analyses: comparison between ages

| Study | Statistical Test | F-Value | P-Value |
| --- | --- | --- | --- |
| PFC ThioS Analysis | Two-Way ANOVA | Interaction F_(2,51)_= 7.629 Time F_(1,51)_= 26.93 Group F_(2,51)_= 15.17 | p=0.0013**  p<0.0001****  p<0.0001**** |
|  | *Bonferroni’s Multiple Comparisons Test* | *2 month WT vs 5 month WT*  *2 month 5XFAD vs 5 month 5XFAD*  *2 month 5XFAD GL-II-73 vs 5 month 5XFAD GL-II-73*  *2 month WT vs 2 month 5XFAD*  *2 month WT vs 2 month 5XFAD GL-II-73*  *2 month 5XFAD vs 2 month 5XFAD GL-II-73*  *5 month WT vs 5 month 5XFAD*  *5 month WT vs 5 month 5XFAD GL-II-73*  *5 month 5XFAD vs 5 month 5XFAD GL-II-73* | p>0.9999  **p<0.0001******  **p=0.012***  p>0.9999  p>0.9999  p>0.9999  **p<0.0001******  **p=0.0005*****  p>0.9999 |
| Hippocampus ThioS Analysis | Two-Way ANOVA | Interaction F_(2,46)_= 7.498 Time F_(1,46)_= 28.69 Group F_(2,46)_= 10.59 | **p=0.0015****  **p<0.0001******  **p=0.0002**** |
|  | *Bonferroni’s Multiple Comparisons Test* | *2 month WT vs 5 month WT*  *2 month 5XFAD vs 5 month 5XFAD*  *2 month 5XFAD GL-II-73 vs 5 month 5XFAD GL-II-73*  *2 month WT vs 2 month 5XFAD*  *2 month WT vs 2 month 5XFAD GL-II-73*  *2 month 5XFAD vs 2 month 5XFAD GL-II-73*  *5 month WT vs 5 month 5XFAD*  *5 month WT vs 5 month 5XFAD GL-II-73*  *5 month 5XFAD vs 5 month 5XFAD GL-II-73* | p>0.9999  **p=0.0001*****  **p=0.0025****  p>0.9999  p>0.9999  p>0.9999  **p<0.0001******  **p=0.0001*****  p>0.9999 |

Table 6. Thioflavin S analyses: comparison between groups

| Study | Statistical Test | F-Value | P-Value |
| --- | --- | --- | --- |
| PFC ThioS Analysis  2months | *Kruskal-Wallis* | *19.79* | **P<0.0001****** |
|  | *Dunn's multiple comparisons test* | *WT Water vs 5xFAD Water*  *WT Water vs 5xFAD GL-II-73*  *5xFAD Water vs 5xFAD GL-II-73* | **P=0.0001*****  **P=0.0022****  p>0.99 |
| Hippocampus ThioS Analysis  2months | *Kruskal-Wallis* | *16.81* | **P=0.0002***** |
|  | *Dunn's multiple comparisons test* | *WT Water vs 5xFAD Water*  *WT Water vs 5xFAD GL-II-73*  *5xFAD Water vs 5xFAD GL-II-73* | **P=0.007****  **P=0.0003*****  p>0.99 |
| PFC ThioS Analysis  5months | *Kruskal-Wallis* | *19.4* | **P<0.0001****** |
|  | *Dunn's multiple comparisons test* | *WT Water vs 5xFAD Water*  *WT Water vs 5xFAD GL-II-73*  *5xFAD Water vs 5xFAD GL-II-73* | **P=0.0003*****  **P=0.0006*****  p>0.99 |
| Hippocampus ThioS Analysis  5months | *Kruskal-Wallis* | *16.81* | **P=0.0002***** |
|  | *Dunn's multiple comparisons test* | *WT Water vs 5xFAD Water*  *WT Water vs 5xFAD GL-II-73*  *5xFAD Water vs 5xFAD GL-II-73* | **P=0.0011****  **P=0.0009*****  p>0.99 |

### ***Supplementary Results and Figures***

#### *Spine count*

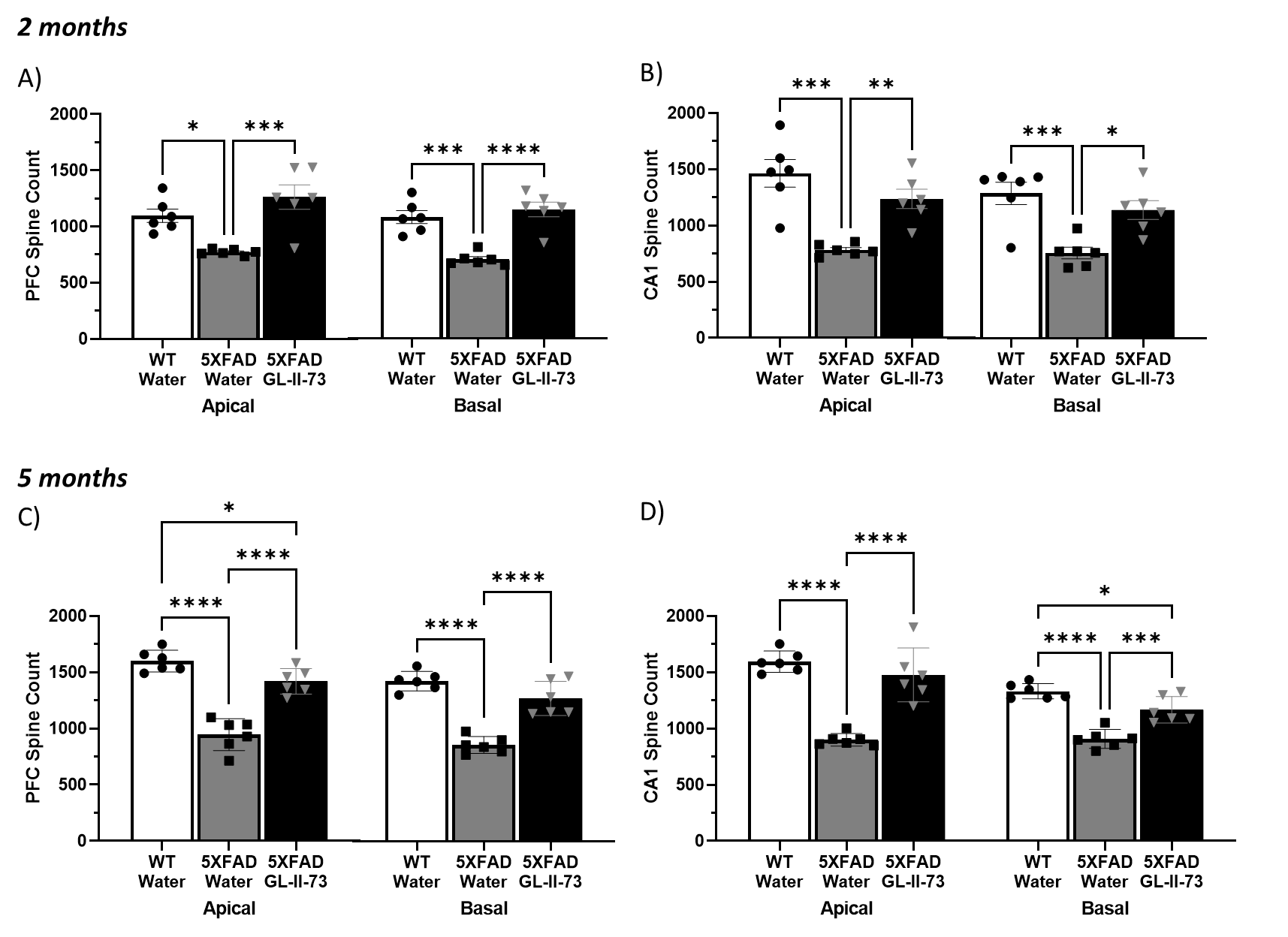

**Supplementary Figure 1.** Effects of GL-II-73 on spine count in the PFC and CA1 of 5xFAD mice, at 2 and 5 months of age

#### *Dendritic length*

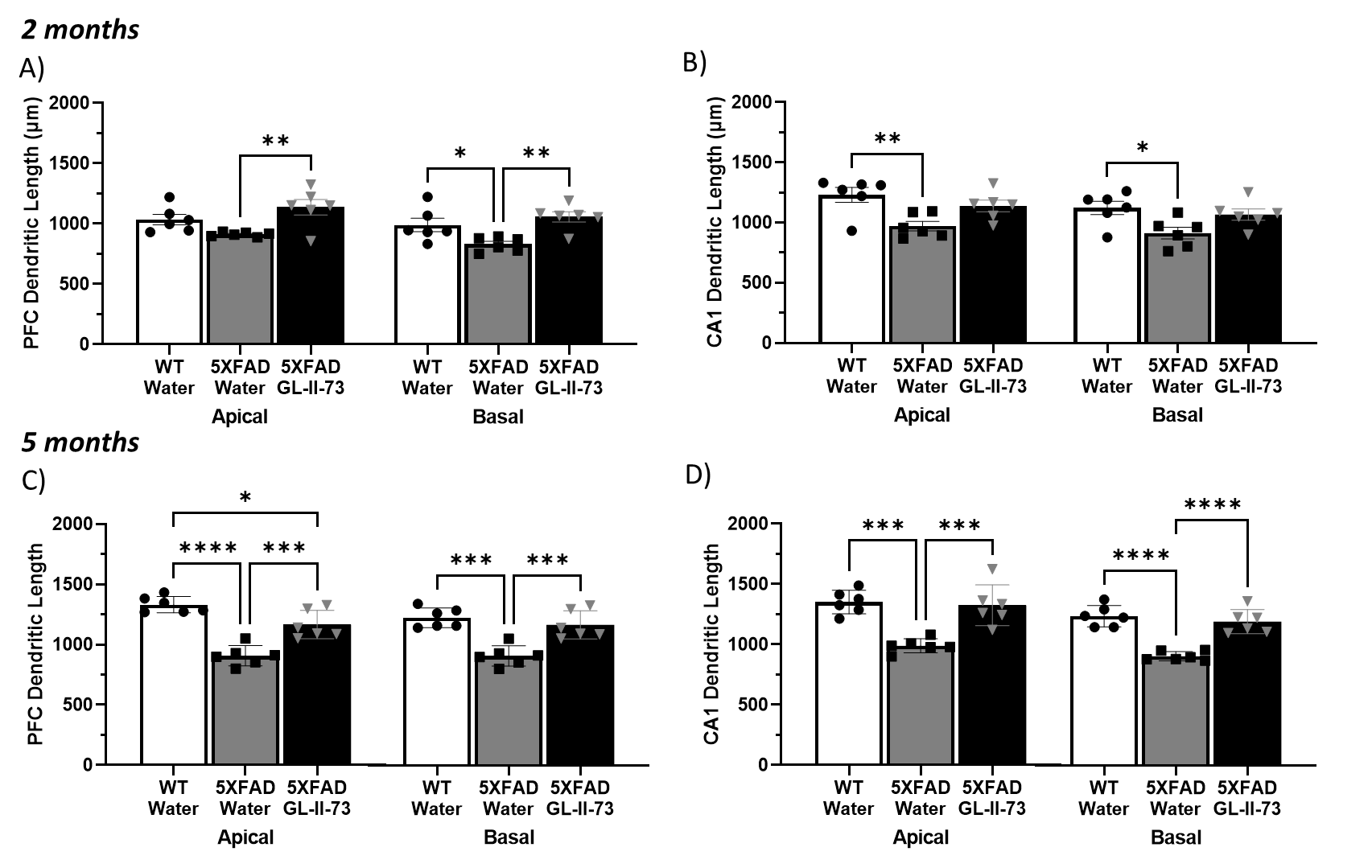

**Supplementary Figure 2.** Effects of GL-II-73 on dendritic length in the PFC and CA1 of 5xFAD mice, at 2 and 5 months of age
